# Not so Apex Predator: Food Web Dynamics in a Freshwater Stream Dominated by Mesopredators

**DOI:** 10.64898/2026.01.30.702877

**Authors:** Celio Roberto Jonck, Jose Marcelo Rocha Aranha

## Abstract

We investigate the impact of apex predator absence on the food web structure and ecological balance of freshwater stream ecosystems at the Salto Morato Natural Reserve, Brazil. Despite the absence of fish due to a natural barrier, species richness in upstream pools was not significantly affected, suggesting ecological balance. Detailed food web analysis revealed that the upstream food web has a higher proportion of top predators, predominantly predatory insect larvae, and lower complexity metrics compared to the downstream web. Our findings challenge the traditional concept of mesopredator release and highlight the unique dynamics of predator-less ecosystems, emphasizing the need for tailored conservation strategies for such fragile environments.

## 1 Introduction

The balance of ecosystems is shaped by predators, whose roles transcend predation itself. Apex predators sustain ecological balance by influencing direct and indirect interactions within food webs (Paine 1966; Natsukawa and Sergio 2022). The importance of apex predators and the significant ecological consequences of their removal can also be seen affecting freshwater ecosystems, such as the loss of biodiversity, water quality, and the ecosystem services they provide (Carpenter et al. 1985; Eby et al. 2006; Vejřík et al. 2017).

Studying freshwater environments in pristine conditions is crucial for understanding natural ecological dynamics and establishing a baseline for ecosystem health. Such knowledge is essential for identifying anthropogenic impacts, developing conservation and management strategies, and comparing the effects of global environmental changes, such as pollution and climate change, enabling more informed and effective responses (Dudgeon et al. 2006; Vörösmarty et al. 2010).

The Salto Morato Natural Reserve, located in the Atlantic Forest of Brazil, offers a unique opportunity to study the dynamics of predator absence in a pristine environment. In this location, we examined the species richness and similarity of pools that belong to the same stream but are separated by a 130-meter-high waterfall. We found that the waterfall acts as a barrier, preventing fish from reaching the upper part of the river (Jonck and Aranha 2010).

The absence of top predators can trigger “mesopredator release.” This phenomenon occurs when top predators are removed from an ecosystem, leading to an increase in the population of mid-level predators. This increase in mesopredators can have significant impacts on the ecosystem, leading to a decrease in biodiversity and changes in the structure of the ecosystem (Ripple and Beschta 2003). Essentially, the removal of top predators disrupts the balance of the food web, leading to cascading effects throughout the ecosystem (Terborgh et al. 2001; Estes et al. 2011).

Conversely, theoretical analyses propose that the absence of an apex predator does not necessarily precipitate ecological instability. Levin (1970) states that no stable equilibrium is possible if some n species are limited by less than n limiting factors. This perspective suggests that equilibrium can be maintained even without an apex predator, provided that there are sufficient limiting factors to balance the ecosystem. Echoing this, McPeek (2014) contended that predator removal might exert minimal impact on consumer richness, especially in scenarios characterized by significant and relatively homogeneous intraspecific density dependence.

This phenomenon was documented in our previous study (Jonck and Aranha 2010), where despite the absence of fish, species richness was not impacted by the presence of the barrier. Our results suggest that the upstream pool, even in the absence of top predators, remains ecologically balanced.

Comprehensive food web analysis is required to understand ecological impacts stemming from the removal of top predators. It helps address ecological problems related to biodiversity loss, pollution, climate change, invasive species, and resource management (Carpenter et al. 1985; Power et al. 1996; Estes et al. 2011; Palkovacs et al. 2011; Marshall et al. 2013; UusiHeikkilä et al. 2018; Price et al. 2019).

Here we aim to analyze the food web properties of the stream community detailed in Jonck and Aranha (2010), focusing on elucidating why the absence of top predators appears to exert no discernible impact on the ecosystem’s equilibrium.

## 2 Material and Methods

Our study was conducted at the Morato River, a third-order stream located at 25° 09’ 54” S and 48° 17’ 50” W. Two pools with similar physical dimensions (approximately 8 m long, 5 m wide, and 3.5 m deep) in the same river flow but separated by a 130 m high waterfall were investigated. Both pool bottoms consisted of a similarly distributed mosaic of sediments, with roughly 60% rocks, 20% gravel, 15% sand, and 5% organic matter accumulations. It is worth noting that the entire drainage basin where the pools are located is within a natural heritage reserve and is not subject to anthropogenic pressures.

Throughout the study, we conducted five sampling campaigns. In each campaign, we collected five samples from each substrate type in both backwaters. We employed reinforced plastic bags measuring 430 x 255 mm as samplers. Sampling involved dragging the bags 10 cm from the surface of sand and gravel substrates and gathering rocks and leaf litter in accordance with the bags’ size. This was executed directly from the substrates using free diving techniques.

Each sample’s material was washed over a 0.05 mm mesh less than two hours from the sampling, and living organisms were sorted using a trans-illuminated tray. The organisms were preserved in 70% alcohol concentration and transported to the laboratory in covered amber glass containers for identification. Each organism found was considered a node in the food web, even if the identification did not reach the species level. The organisms were identified to the lowest possible taxonomic level, and distinct but unresolved species were considered morphospecies. Fish species presence was assessed via visual census, conducting two one-hour assessments per day - one in the morning and another at dusk - during each campaign, using Barreto and Aranha (2005) as a guide for species identification.

The diet of organisms in the backwater’s communities upstream and downstream of the waterfall was established based on specialized literature references (Merritt and Cummins 1996a; Merritt and Cummins 1996b; Motta and Uieda 2004; Motta and Uieda 2005; Aranha 2000; Di-Sabatino et al. 2000; Dole-Oliver et al. 2000; Albertoni et al. 2003; Jaarsma et al. 1998; Gibran and Souza 2001; Kehl and Dettner 2003; Rosi-Marshall and Wallace 2002; Chivers and Mirza 2000; Ranvestel et al. 2004; Aranha et al. 1998; Aranha et al. 2000; Henriques-Oliveira et al. 2003). A spatial criterion was also employed, confirming a trophic link only if both consumer and resource were found on the same substrate. Highly mobile organisms were considered occupants of all substrates. The trophic web was mapped using a binary matrix approach (Cohen et al. 1990).

Basal nodes were categorized according to the consumed resource type, independent of their taxonomic composition. Consequently, microbial biofilms on rocks and organic substrates, irrespective of the microbial diversity, were associated with scraper feeding activities. Similarly, algae and bryophytes were associated with grazers, suspended organic matter was associated with filter feeders, plant detritus was associated with shredders, and animal detritus was associated with scavengers.

Ten properties of the backwaters’ webs at both locations were analyzed to infer the waterfall’s influence. These properties included the proportions of top, intermediate, basal, and omnivorous species, calculated by dividing the number of species in each category by the total number of species. The number of mesopredators, calculated as the number of nodes that feed on a consumer, are not consumed by other nodes but are not at the top trophic level of the food web. The maximum and the minimum length of the web was noted as the largest and the smallest number of links between a basal and a top species. The density of links in the web was assessed by dividing the total number of links by the number of species, connectance was measured as the total number of links divided by the total number of mathematically possible links in the web, and the number of trophic compartments was defined as isolated sectors in the web containing representatives of all trophic levels but with no connections between their elements.

## 3 Results

In the backwater located upstream of Salto Morato, the constructed food web comprises 76 nodes. Among these, five basal nodes were categorized based on consumer feeding behaviors. The remaining 71 nodes correspond to organisms collected from various substrates within the backwater area. Predominantly, these organisms were aquatic insects, with notable representations from the Trichoptera (16 taxa), Coleoptera (12 taxa), and Ephemeroptera (10 taxa) groups. A significant observation was that a majority of these organisms (45.8%) were exclusive to a single substrate, whereas a mere 6.9% spanned across all considered substrates (Table SM1).

The food web of the backwater located downstream of Salto Morato was constructed with 70 nodes. The same five basal nodes plus 65 representing organisms collected across different substrates. Here, over half of the organisms (50.8%) were identified in unique substrates, and 20.0% of taxa were found across all substrates. The taxonomic composition majorly included aquatic insects, with a noteworthy addition of fish taxa (12 taxa). Ephemeroptera emerged as the most populous insect group (11 taxa), alongside a significant representation of mites (Actinedida) with 9 taxa (Table SM2).

Analysis indicates substantial structural differences in the upstreams food web (Figure 1) when compared with the downstream food web (Figure 2).

**Figure 1.**
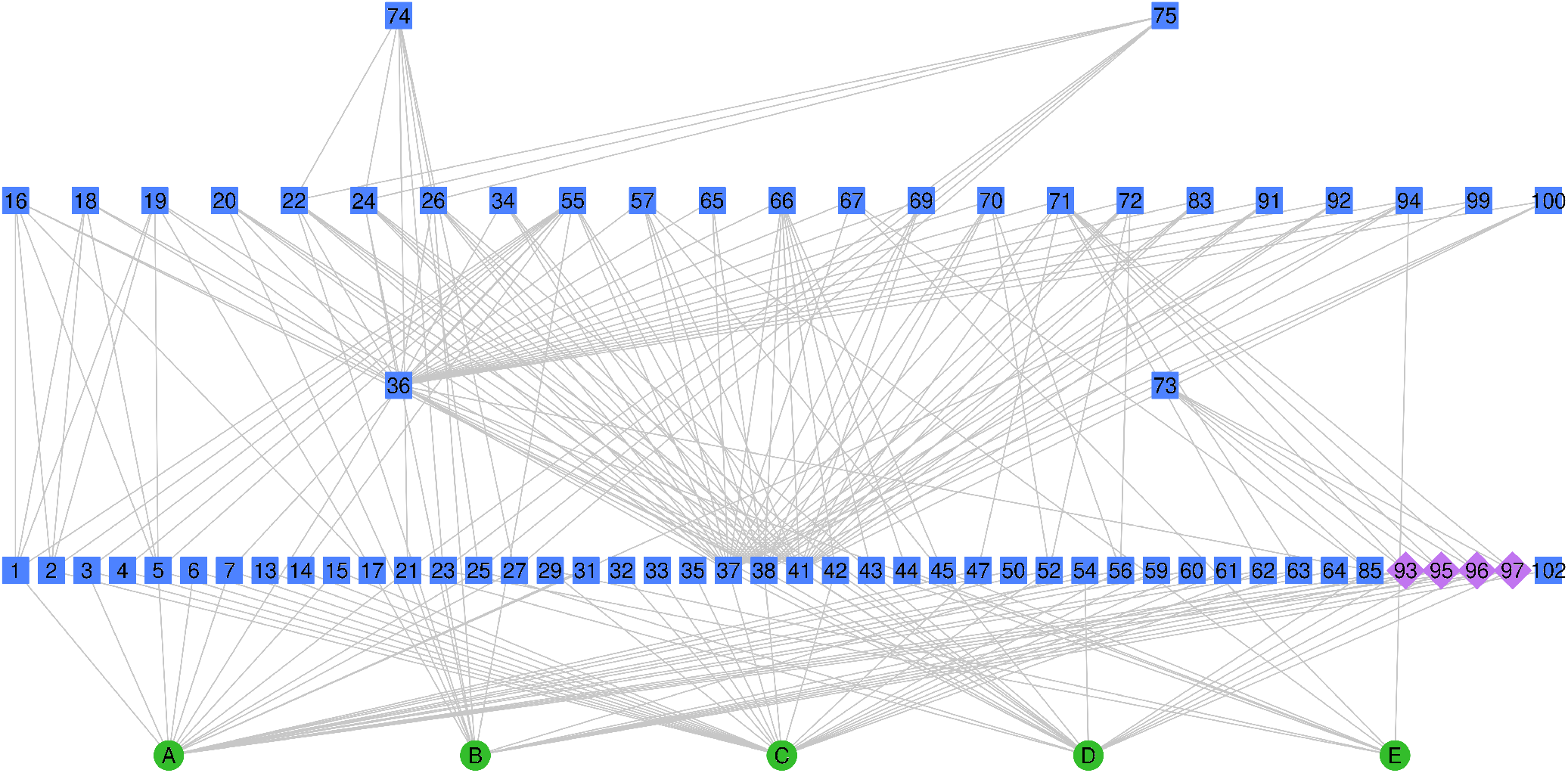
Food web of the upstream pool. A-Biofilm, B-Algae and Bryophytes, C-Particulate Organic Matter, D-Vegetal Detritus, E-Animal Detritus, 001-Paracloeodes, 002-Cloeodes, 003-Askola, 004-Tricorytopsis, 005-Apobaetis, 006-Farrodes, 007-Miroculis, 013-Campylocia, 014-Leptohyphodes, 015-Traulodes, 016-Anacroneuria, 017-Tupiperla, 018-Macroginoplax, 019-Kempnya, 020-Triplectides, 021-Phylloicos, 022-Grumichella, 023-Oecetis, 024-Nectopsyche sp1, 025-Calamoceratidae sp1, 026-Nectopsyche sp2, 027-Helicopsyche, 029-Cernotina, 031-Glossosomatidae sp1, 032-Polyplectropus, 033-Austrotinodes, 034-Marilia, 035-Leptonema, 036-Tanypodinae, 037-Chironominae, 038-Orthocladiinae, 041-Stenochironomus, 042-Tipulidae sp1, 043-Ceratopogon, 044-Coulicoides, 045-Monohelea, 047-Stenelmis, 050-Psephenus, 052-Hexacylloepus, 054-Hydrophilidae sp1, 055-Dytiscidae sp1, 056-Ordobrevia, 057-Gyretes, 059-Neoelmis, 060-Promoresia, 061-Adult Hydrophilidae, 062-Macrelmis, 063-Phanoceros, 064-Lutrochus, 065-Limnochoris, 066-Pleochoris, 067-Cryphocricos, 068-Paraplea, 069-Hetaerina, 070-Limnetron, 071-Brechmorhoga, 072-Elga, 073-Neocordulia, 074-Progomphus, 075-Epigomphus, 083-Hydrachnidia sp5, 085-Naididae, 091-Hydrachnidia sp10, 092-Hydrachnidia sp11, 093-Anura sp1, 094-Dugesidae, 095-Anura sp2, 096-Anura sp3, 097-Anura sp4, 099-Hydrachnidia sp12, 100-Hydrachnidia sp13, 102-Dryopidae sp2.

**Figure 2.**
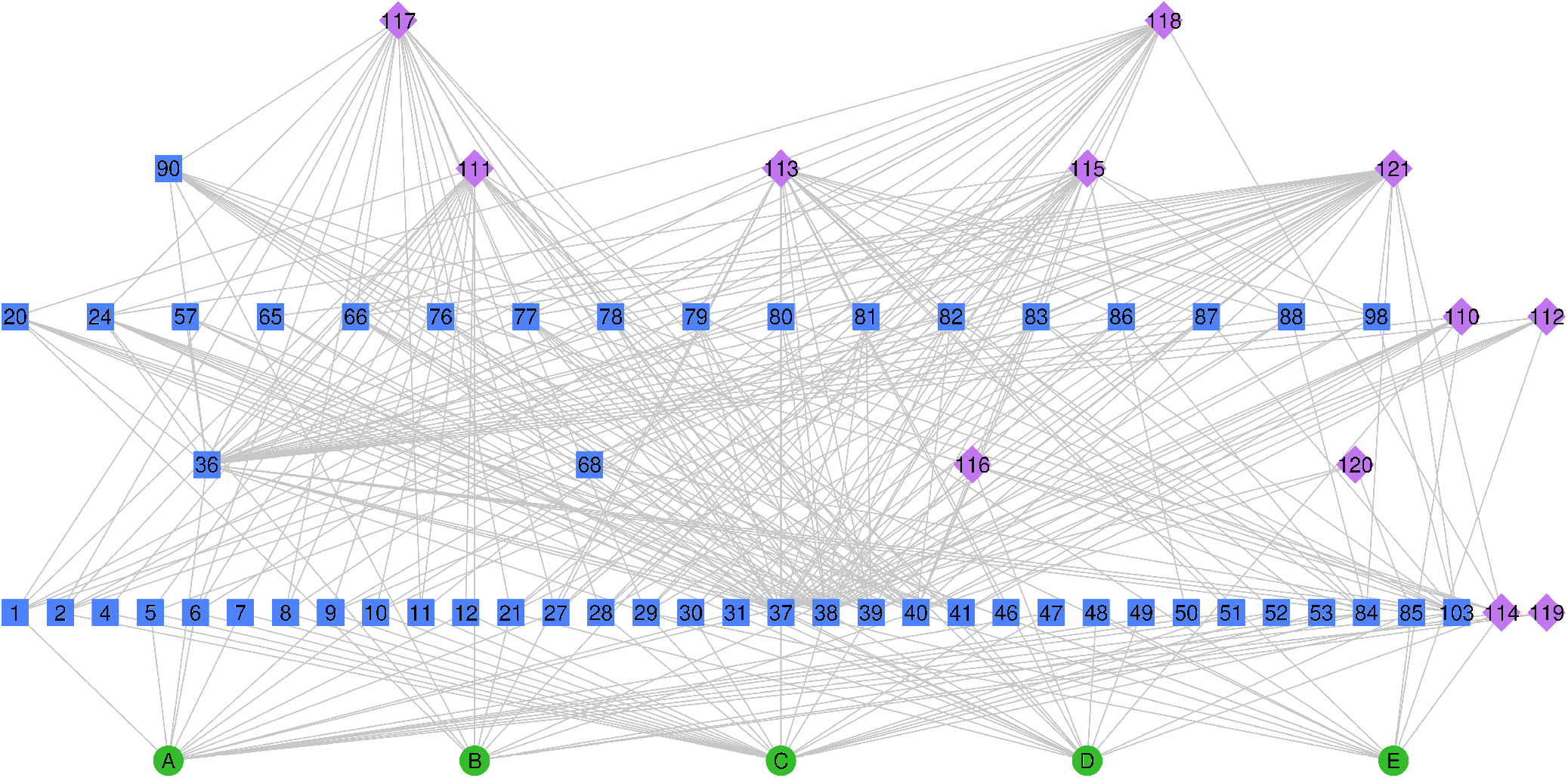
Food web of the downstream pool. A-Biofilm, B-Algae and Bryophytes, C-Particulate Organic Matter, D-Vegetal Detritus, E-Animal Detritus, 001-Paracloeodes, 002-Cloeodes, 004-Tricorytopsis, 005-Apobaetis, 006-Farrodes, 007-Miroculis, 008-Traverhyphes, 009-Aturbina, 010-Americabaetis, 011-Tricorythodes, 012-Hylister, 020-Triplectides, 021-Phylloicos, 024-Nectopsyche sp1, 027-Helicopsyche, 028-Macronema, 029-Cernotina, 030-Chimarra, 031-Glossosomatidae, 036-Tanypodinae, 037-Chironominae, 038-Orthocladiinae, 039-Simulidae, 040-Bezzia, 041-Stenochironomus, 046-Heterelmis, 047-Stenelmis, 048-Polyphaga 1, 049-Scolytidae, 050-Psephenus, 051-Xenelmis, 052-Hexacylloepus, 053-Xenelmis, 057-Gyretes, 065-Limnochoris, 066-Pleochoris, 068-Paraplea, 076-Oxystigma, 077-Heteragrion, 078-Argia, 079-Hydrachnidia sp1, 080-Hydrachnidia sp2, 081-Hydrachnidia sp3, 082-Hydrachnidia sp4, 083-Hydrachnidia sp5, 084-Cladocera, 085-Naididae, 086-Hydrachnidia sp6, 087-Hydrachnidia sp7, 088-Hydrachnidia sp8, 090-Macrobrachium, 098-Hydrachnidia sp12, 103-Copepoda, 110-Astyanax, 111-Deuterodon, 112-Mimagoniates, 113-Corydoras, 114-Phalloceros, 115-Characidium, 116-Geophagus, 117-Rhamdia, 118-Pimelodella, 119-Rineloricaria, 120-Ancistrus, 121-Crenicichla.

The downstream web is characterized by a classical diamond configuration, wherein the bulk of nodes is located at the intermediate trophic levels. Conversely, the upstream web is delineated by an inverted pyramid structure, indicating a concentration of nodes at higher trophic levels. Despite the physical compartmentalization of the environment, no compartmentalization of the food web was observed in any of the studied backwaters.

It’s noteworthy that, although the upstream web encompasses a greater total number of trophic nodes, it comprises one fewer trophic level compared to its downstream counterpart. This observation extends to the measures of omnivory, trophic links, linkage density, and connectance, all of which are quantitatively lower in the upstream web. However, the relative abundance of mesopredators in the upstream web is nearly triple the number found in the downstream web. Detailed results are presented in Table 1.

**Table 1.** Comparison of Food Web Properties.

|  | Upstream |  | Downstream |  |
| --- | --- | --- | --- | --- |
|  | n | % | n | % |
| Trophic nodes | 76 |  | 70 |  |
| Basic nodes | 5 | 6.5 | 5 | 7.1 |
| Intermediate nodes | 33 | 42.9 | 51 | 72.9 |
| Top nodes | 39 | 50.6 | 14 | 20 |
| Omnivores | 26 | 33.8 | 31 | 44.3 |
| Mesopredators | 21 | 27.3 | 8 | 11.4 |
| Trophic levels | 5 |  | 6 |  |
| Maximum length | 4 |  | 5 |  |
| Minimum length | 1 |  | 1 |  |
| Trophic compartments | 0 |  | 0 |  |
| Trophic links | 226 |  | 324 |  |
| Link density | 2.94 |  | 4.63 |  |
| Connectance | 0.04 |  | 0.07 |  |

## 4 Discussion

The food webs were constructed with 76 nodes upstream and 70 nodes downstream. Since we standardized the available resources in the streams into five categories, the number of nodes indicates the species richness we found for each pool: 71 upstream and 65 downstream. This number can vary considerably with sampling effort, but studies in similar lotic environments found richness values consistent with those presented here (Melo and Froehlich 2001; Clarke et al. 2008; Bacca et al. 2023). While we acknowledge that our sampling is limited and that greater sampling effort could alter the characteristics of the food webs (Goldwasser and Roughgarden 1997; Pringle and Hutchinson 2020), we are confident that the communities of both pools are represented in our sampling and that cryptic species, if they exist, would not resolve the identified discrepancies.

The proportion of basal, intermediate, and top species in the downstream food web is 0.071, 0.729, and 0.2, which aligns with the proportional ranges of medium-resolution webs compiled by Schmid-Araya et al. (2002b). Although these proportions do not perfectly match the ranges presented by the author for high taxonomic resolution webs (Schmid-Araya et al. 2002b; Schmid-Araya et al. 2002a), the web’s structure follows the same pattern, with a higher proportion of intermediate species than basal or top species. These data indicate that the downstream web of Salto Morato can be considered a regular web.

Conversely, the upstream web structure at Salto Morato differs significantly, with proportions of 0.065 basal species, 0.429 intermediate species, and 0.506 top species. This characteristic, a higher proportion of top species than intermediate species, has never been documented in balanced ecosystems, appearing only as a transitional condition in manipulated experiments (Estes and Palmisano 1974; Babcock et al. 1999; Wilkinson et al. 2022) or unbalanced environments (Hughes et al. 2007; Harvey and MacDougall 2015; Essert et al. 2024).

All properties of a food web usually associated with complexity and stability, such as trophic levels, omnivory, trophic links, linkage density, and connectance (Warren 1990; McCann and Hastings 1997; Melián and Bascompte 2004; Scotti et al. 2009; Kratina et al. 2012; Awender et al. 2021; Vitekere et al. 2021), are lower in the upstream web. The only property of the upstream web that exceeds that of the downstream web is the proportion of mesopredators, whose release is associated with imbalances caused by the removal or loss of top predators (Ripple and Beschta 2003).

These results led us to question how such an unbalanced food web could be associated with an almost pristine environment. The answer lies in the linkage density and reevaluation of the mesopredator concept we used.

Since the two pools are located in the same stream and have the same substrate types, we can infer from the condition of the downstream web that there is no energy limitation for the upstream web. Therefore, the expected linkage density for these environments would be the same, but it is reduced in the upstream web to two-thirds of the downstream web value. However, when comparing the three species with the highest linkage densities in the two webs, we see that Tanypodinae has 30 links upstream and 34 downstream, Chironominae has 23 links upstream and 29 downstream, and Orthocladiinae has 22 links upstream and 29 downstream. Thus, the upstream web still has a lower value, but the difference is much smaller, indicating that these species form a cohesive cluster on which the rest of the web depends. Supporting this assertion, we see that almost all organisms in higher trophic levels, many of which were identified as mesopredators, use these three species as a resource.

Moreover, we need to revise the concept of mesopredators to understand what happens in the pools. The concept of mesopredators was introduced in the study by Ripple and Beschta (2003), which investigated the ecological role of wolves and the ecological consequences of their removal from their ecosystem. The mesopredator release refers to the phenomenon where the removal or decline of top predators led to an increase in medium-sized predator populations (mesopredators), such as coyotes and raccoons. This changed the ecosystem dynamics with a cascading effect, as large herbivores, which were preyed upon by wolves but not by mesopredators, thrived and imposed greater pressure on vegetation. The lack of vegetation caused a decrease in food and shelter for other species and even soil erosion, affecting the entire ecosystem.

As elegant as the concept is, this does not seem to be the case for the upstream food web of Salto Morato. The dynamics of the Morato River indicate that the fish, which are the top predators of the downstream pool, did not get removed from the upstream pool but never had access to it due to the barrier imposed by the waterfall (Jonck and Aranha 2010). We know that several species of predatory insect larvae would be prey in an environment with fish, but in the case of the upstream pool of the Morato River, they are not mesopredators released by the occurrence of an imbalance; they are the top predators of the ecosystem.

Resolving this does not make the upstream web a regular web, obviously, but it removes the bias of thinking that everything outside the regular is unbalanced or altered and allows us to conclude that the lotic ecosystem above the waterfall is a very fragile environment and that its conservation or the restoration of similar degraded environments can be very challenging.

## 5 Conclusion

We examined the impact of apex predator absence on the food web structure and ecological balance in freshwater streams at the Salto Morato Natural Reserve. Detailed food web analysis revealed significant structural differences between the environments, with the upstream pool characterized by a higher proportion of mesopredators and lower complexity metrics.

These findings challenge the traditional concept of mesopredator release, suggesting that ecosystems can develop unique dynamics in the absence of apex predators. This underscores the importance of considering local environmental conditions and historical development when assessing ecosystem stability and resilience.

Our study highlights the need for tailored conservation strategies for fragile environments like the upstream pool of the Morato River. Future research should focus on exploring the long-term stability of these ecosystems in a world that faces climate changes and increasingly intense anthropogenic pressures.

By providing insights into the unique dynamics of this mesopredator dominated ecosystem, this study contributes to a broader understanding of food web structures and their role in sustaining biodiversity and ecological services. Our findings emphasize the complexity of ecological interactions and the necessity for comprehensive approaches in ecosystem management and conservation efforts.

---

**Table SM1.**
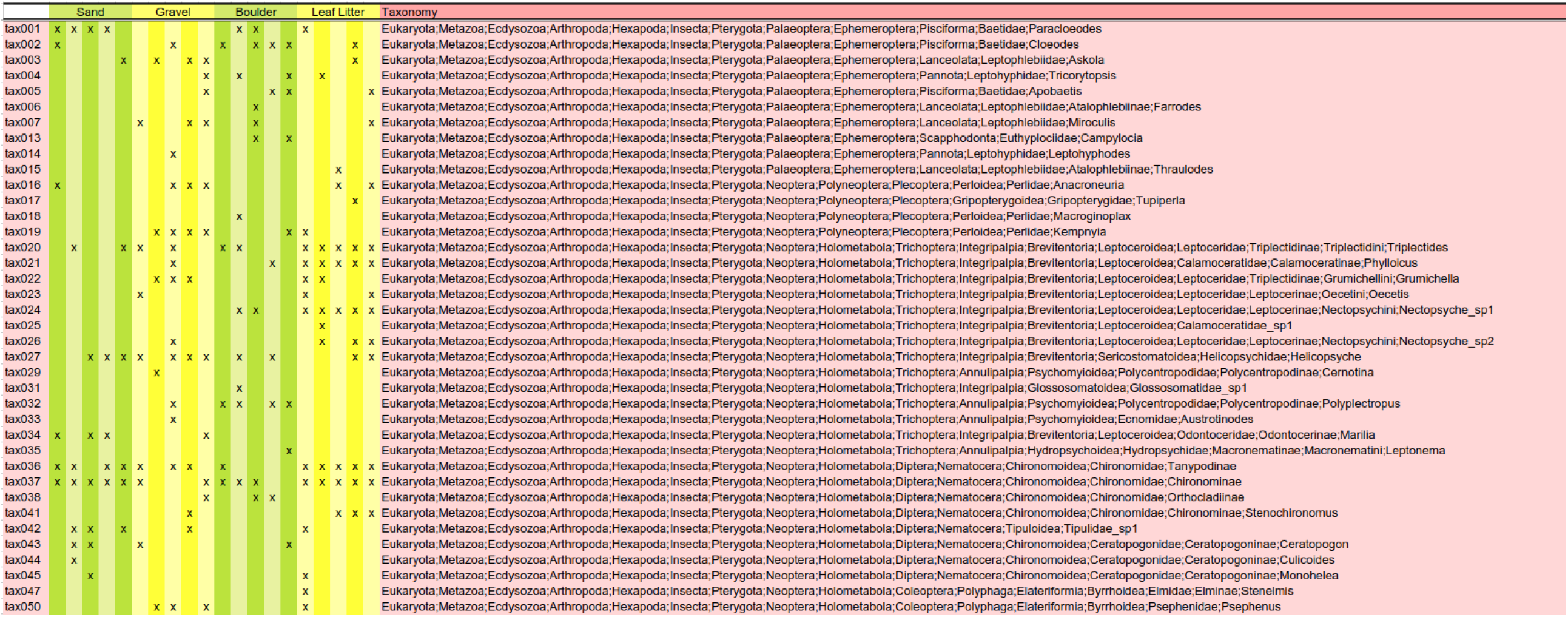
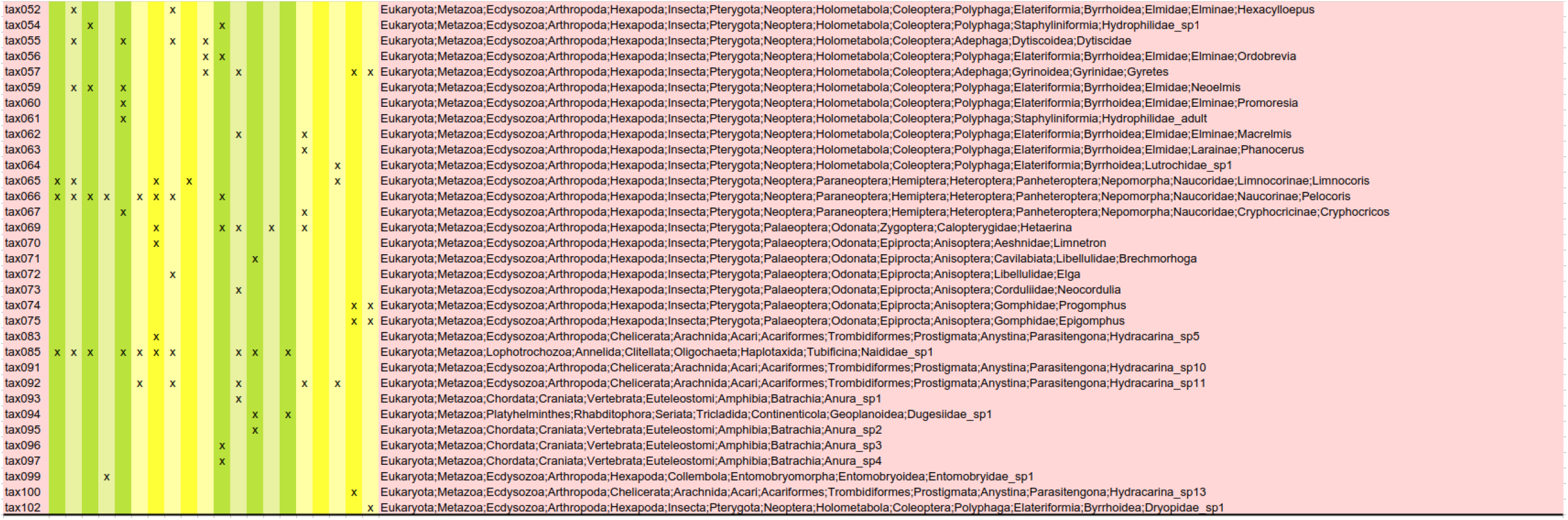

**Table SM2.**
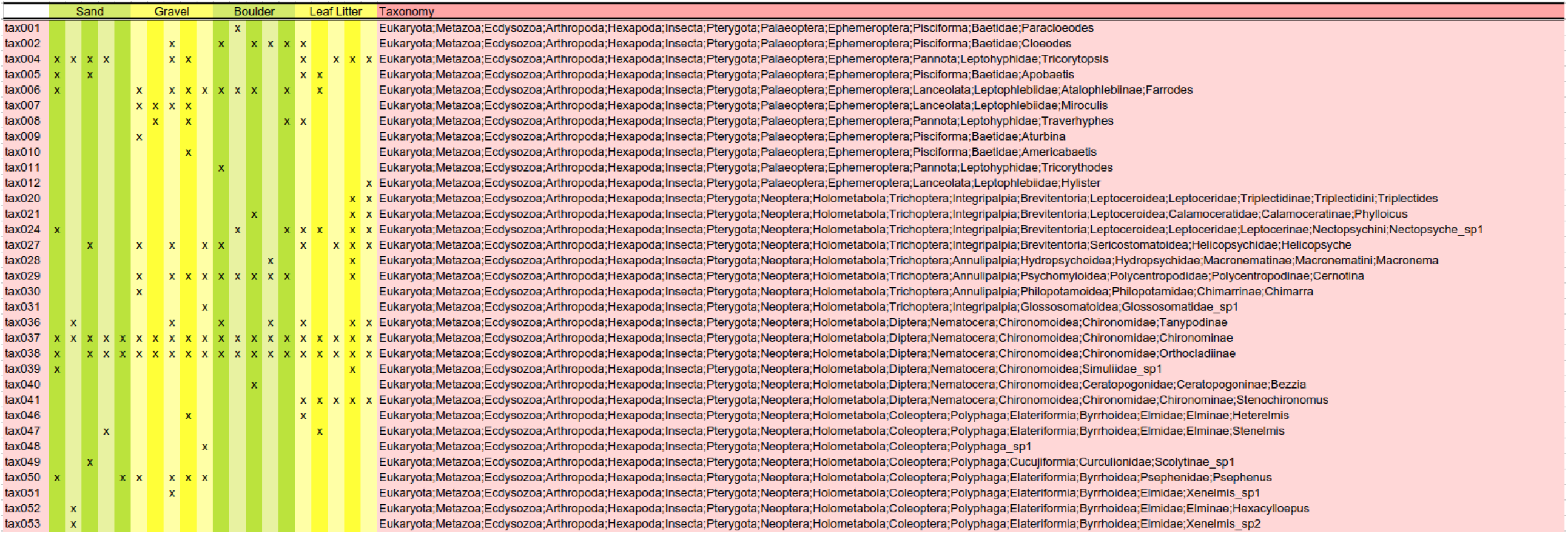
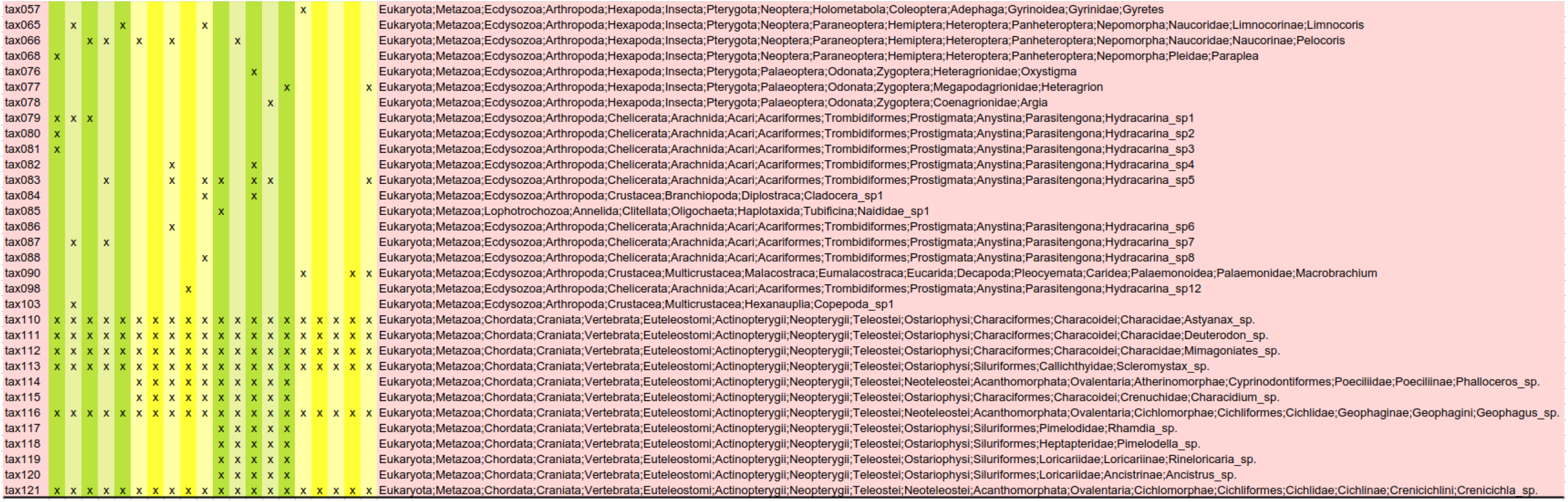

